# Scaling temperature-dependent dispersal rates to metacommunity dynamics: An experimental test

**DOI:** 10.64898/2026.05.21.727003

**Authors:** Keila Stark, Ze-Yi Han, Jean P. Gibert, Mary I. O’Connor

**Affiliations:** Kellogg Biological Station, Michigan State University, Hickory Corners, USA; Department of Biology, Duke University, Durham, USA; Biodiversity Research Centre and Department of Zoology, University of British Columbia, Vancouver, Canada

**Keywords:** protist, metabolic theory of ecology, thermal response curve, *Paramecium*, *Tetrahymena*, competition-colonization trade-off

## Abstract

1. Changes in community structure under shifting thermal regimes depend on how both local population dynamics and regional dispersal respond to temperature. Processes underlying dispersal, such as movement speed and density dependence, are constrained by temperature-dependent metabolic rates; however, the temperature dependence of population dispersal rate, and effect of this relationship on local and regional diversity patterns, have received little attention in the metabolic scaling literature.
2. Here, we propose and experimentally test a framework that relates temperature effects on individual dispersal probability, to thermal performance curves (TPCs) for population dispersal rates, to colonization dynamics in metacommunities. Using multi-patch well plate microcosms, we measured thermal performance curves for dispersal rate in several naturally co-occurring ciliate species, and contrasted species-specific dispersal TPCs at different intra- and inter-specific densities and time scales.
3. Dispersal rate TPCs in monoculture differed at low versus high population densities, potentially suggesting distinct temperature effects on the density-independent (individual movement probability and speed) and density-dependent (quorum-sensing and resource competition) components of dispersal.
4. Species-specific dispersal rate TPCs in polyculture metacommunities explained differences in colonization dynamics across temperature treatments. Dispersal rate TPCs differed from intrinsic growth rate TPCs, such that better dispersers had higher-than-expected per capita population growth at the regional (whole-metacommunity) scale compared to predictions from standard growth TPCs measured in single-patch monoculture.
5. Together, these results suggest that ignoring temperature-dependent dispersal can yield an incomplete understanding of biodiversity change in spatially structured systems exposed to warming.

## 1 Introduction

Dispersal shapes patterns of genetic, phenotypic, and taxonomic diversity across spatially structured landscapes (Cadotte, 2006; Cote et al., 2017; Loreau et al., 2003; Ronce and Clobert, 2012; Suárez et al., 2022; Yamaguchi, 2022). In the context of global change, dispersal can facilitate community resilience against environmental perturbations at local scales (Loreau et al., 2003; Mouquet and Loreau, 2003) and rates of range expansion at regional scales (Norberg et al., 2012; Williams and Blois, 2018). Experiments in which dispersal was manipulated by the researchers suggest that dispersal magnitude mediates changes in community structure in systems exposed to warming (Grainger and Gilbert, 2017; Parain et al., 2019; Thompson et al., 2015). Less appreciated is the fact that temperature can also affect population dispersal rates themselves, affecting abundance and diversity separately from temperature-dependent population growth rates (but see Amarasekare 2024). Metabolic scaling theories explain how biological rates that underlie community dynamics respond to warming based on the metabolic temperature dependence of processes occurring at lower levels of biological organization (O’Connor et al., 2011; Rall et al., 2009; Wieczynski et al., 2021). For example, temperature-dependent velocity in individuals has been scaled up to derive temperature-dependent consumer-resource encounter rates (Gibert et al., 2016; Pawar et al., 2012). At the population level, unimodal thermal performance curves (TPCs) for intrinsic growth rates are well-documented and understood (Eppley, 1972; Savage et al., 2004; Thomas et al., 2012), but this is not so for other population-level processes influencing community structure. A lack of theory and data on TPCs for population dispersal rates, defined as the net proportion of a population moving between two patches per unit time (ind ind^-1^ time^-1^), hinders a general understanding of how local and regional diversity patterns respond to warming (Amarasekare, 2024).

In motile and sessile organisms alike, population dispersal generally consists of three phases (Clobert et al., 2009; Travis et al., 2013): (1) Emigration, where individuals or propagules leave their natal habitat; (2) Transport, driven by passive forces like wind and currents, or active locomotion; and (3) Settlement and recruitment, where individuals arrive in a suitable habitat patch and reproduce. This framework suggests that dispersal rates depend not only on transport rates but also on demographic density dependence and reproduction. These multiple sources of variability appear to preclude a generalized model of temperature-dependent dispersal that can be applied across systems; however, temperature-dependent metabolism—a feature of all living things—constrains temperature’s effects on all three phases of dispersal through locomotion (Gibert et al., 2016), development rates (Gillooly et al., 2002), population growth (Savage et al., 2004; Thomas et al., 2012), and density dependence (Lemoine, 2019; Vasseur, 2020). Energy produced via aerobic respiration allows contractions in locomotor cells and organelles (e.g., muscle cells, cilia) to propel motile organisms, such that variation in movement rates across broad taxonomic groups, including routine velocity, attack and escape velocity, and angular momentum is explained by allometric body size scaling and temperature-dependent enzyme kinetics that are central to metabolic scaling theory (Hurlbert et al., 2008; McNab, 1963; Pawar et al., 2012; Peters, 1983; Ritchie et al., 2001; Schmidt-Nielsen, 1972; Sleigh, 1956; Taylor, 1963; Tsubaki et al., 2010). The metabolic mass dependence of velocity has been used to explain variation in movement occurring at broader spatiotemporal scales, such as home range movements and migratory distances (Doughty et al., 2016; Hein et al., 2012; Hillaert et al., 2018; Jetz et al., 2004; Tamburello et al., 2015). With regard to temperature, few studies have used metabolic scaling to link the temperature dependence of instantaneous organism movements to those at broader spatiotemporal scales and levels of organization. Understanding the temperature scaling of population dispersal rates remains an important knowledge gap due to its implications for demography, community assembly, and range dynamics under global change.

The temperature dependence of population dispersal may be understood by scaling up from individual movement rates, to dispersal probabilities, to population dispersal rates, and finally, to colonization rates in metacommunities. Each step upwards in scale has theoretical and empirical gaps. First, metabolic theory does not predict how individual movement rates translate to the probability of dispersal between habitat patches. Even when instantaneous movement speeds increase with temperature, this does not necessarily translate to an increased likelihood of an individual dispersing to an adjacent habitat patch (Barnes et al., 2015) due to genetic, phenotypic, and behavioural factors not captured by metabolic theory’s simple canonical equations (Bestion et al., 2015; Jacob et al., 2018). Second, it is unclear whether temperature effects on individual dispersal probability directly scale to effects on population dispersal rate in organisms experiencing density dependence. Resource-mediated density dependence changes with temperature in ectothermic taxa via changes in consumption rates driven by metabolic demands when resource supply is constant (Bernhardt et al., 2018); density dependence may thus affect dispersal TPCs by modifying per capita emigration rates through quorum-sensing or direct resource competition (Matthysen, 2005; Travis et al., 1999). This effect occurs at all temperatures but is likely strongest at those where competition is strongest and resource availability is lowest (which in turn vary depending on thermal response curves for birth rates, death rates, and resource availability; Amarasekare 2015; Bernhardt et al. 2018; DeLong and Hanson 2009; Lemoine 2019; Vasseur 2020). Finally, thermal response curves for intrinsic population dispersal rates measured in monoculture populations may vary from realized thermal response curves for dispersal in a metacommunity context with multiple interacting species. Interspecific variation in dispersal TPCs in multi-species contexts may reflect correlations or trade-offs in species’ dispersal and growth abilities across temperatures. For example, if a species with poor growth at a given temperature is a good disperser at that temperature (i.e., competition-colonization trade-off), then in a metacommunity context it may persist even if it would go locally extinct based on its thermal performance curve for local population growth.

We systematically tested for similarities and differences in thermal response curves for dispersal across levels of organization in a series of experiments using six ciliate species in multi-patch microcosms. We tested three hypotheses to tease apart temperature effects on the different components of population dispersal: (H1) Dispersal rates are temperature-dependent at both low (1a) and high (1b) population densities; (H2) The strength of temperature effects on dispersal differs at low versus high population densities, reflecting distinct effects on individual movement and population density dependence; (H3) Interspecific variation in dispersal thermal performance curves explain differences in metacommunity dynamics and diversity patterns across temperatures.

## 2 Materials and Methods

### 2.1 Study system and experimental setup

To investigate the temperature dependence of population dispersal rates, we conducted dispersal experiments on six freshwater ciliate species in monoculture: *Paramecium caudatum*, *Paramecium aurelia*, *Paramecium bursaria*, *Colpidium striatum*, *Glaucoma* sp., and *Tetrahymena pyriformis*. Ciliates are an ideal system for testing our hypotheses (Altermatt et al., 2015) because they are motile and spatially structured metacommunities can be easily constructed in laboratory settings (Arancibia and Morin, 2022; Jacob et al., 2018). Their short generation times allow short-term dispersal to be measured over minutes and longer-term dispersal and population growth to be measured over days. All species are bacterivores within the same trophic level, so they likely are in competition and there is no within-community predation. Protist cultures were obtained from Carolina Biological Supply (Burlington, NC, USA) and maintained at Duke University before transport to the University of British Columbia where the experiments were conducted. Stock cultures were maintained in 250 mL glass Erlenmeyer flasks at 22 °C under a 12-hour light:dark cycle in Timothy hay media. Protists were fed communities consisting of thousands of bacteria collected from an ephemeral pond at Duke Forest (Wilbur/Gate 9 pond: 36.013914, -78.979720) as described in Rocca et al. (2022). We added an autoclaved wheat seed to each culture as a carbon source for bacteria. Three days before each experiment, stock cultures were replenished with fresh media and transferred to a dark regime to remove photosynthetic endosymbionts of *Paramecium bursaria*.

All experiments were conducted in 24-well plates filled with 1 mL of hay protist media, where wells (patches) were connected by media-filled glass capillary tubes bent at a 90-degree angle over a flame following Arancibia and Morin (2022). The protist media in the capillary tubes was sterile (autoclaved). Media in the wells (patches) was taken from a single stock polyculture and vacuum filtered through polycarbonate filter paper (0.8 *μm* pore size) to remove all protists while retaining their shared bacterial food source. Replicates in all three experiments were repeated at the same six temperatures (14 °C, 18 °C, 22 °C, 26 °C, 30 °C, 34 °C). This temperature range was chosen to capture all species’ full thermal range for population growth (Wieczynski et al., 2021).

To distinguish temperature effects on the density-independent and density-dependent components of dispersal (H1 and H2), we performed identical short-term dispersal assays at low (Experiment 1) and high population densities (Experiment 2). To do so, we used monocultures of each species in two-patch arenas, where one well served as the inoculation patch and the other served as the empty recipient patch (Figure 1). To test whether differences in dispersal TPCs contribute to temperature-associated differences in regional dynamics and metacommunity structure (H3), we conducted a multi-patch colonization experiment using polycultures of five species that were easy to visually distinguish (Experiment 3).

**Figure 1:**
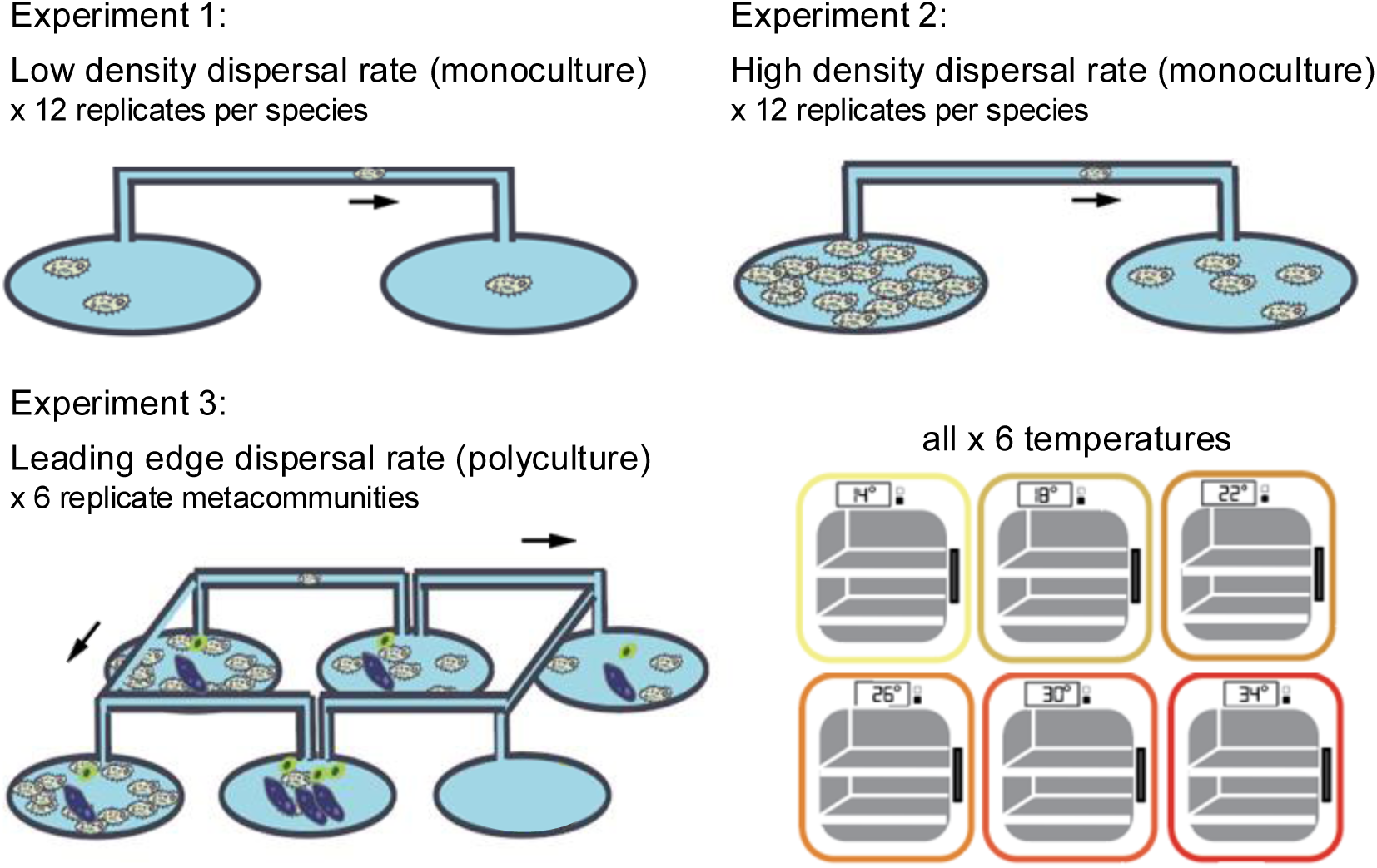
Conceptual diagram showing microcosm arrangements for the three dispersal experiments. Dispersal at low (Experiment 1) and high population densities (Experiment 2) was assayed using monocultures of each species in two-patch arenas. Leading edge dispersal rate, or dispersal rates underlying metacommunity colonization, were quantified in ring-shaped multi-patch microcosms containing polycultures. Replicate numbers are per temperature treatment: each assay was replicated all 6 temperatures, including each species in the monoculture experiments 1 and 2.

### 2.2 Experiment 1: Low-density dispersal in monoculture

For the low density dispersal assays, the source patch was inoculated with 5 ± 1 individuals in assays using the *Paramecium* species, or a roughly equivalent biovolume of the smaller species (Figure 1, Table S2). The assay was replicated 12 times in each species at each treatment temperature (Table 1). After inoculation, we placed well plates in incubators set to their respective treatment temperatures for 300 minutes. We checked them every 15 minutes until a dispersal event was observed and the time until first dispersal (min) was recorded for the replicate. Replicates in which no dispersal event was observed by the end were assigned a wait time of 300 minutes and retained in the analysis. Low density dispersal rates were calculated as 1/*N_i_*/*t*, where *N_i_* is the species-specific inoculation density (see Table S2) and *t* is time until first dispersal event in minutes. A preliminary pilot run of this assay confirmed that back-dispersal into the inoculation patch was unlikely within the assay duration.

**Table 1:** Scale of inference and number of replicates for processes of interest. Note that the number of replicates corresponds with one single temperature treatment; the full replication regime described in this table was repeated at six different temperatures (14 °C, 18 °C, 22 °C, 26 °C, 30 °C, 34 °C).

| Process of interest | Experiment | Scale of inference | Scale at which the factor of interest is applied | Number of replicates |
| --- | --- | --- | --- | --- |
| Dispersal rate, low-density monoculture | 1 | Population | 2-patch microcosm | 12 |
| Dispersal rate, high-density monoculture | 2 | Population | 2-patch microcosm | 12 |
| Dispersal rate, polyculture | 3 | Population | 7-patch microcosm | 6 |

### 2.3 Experiment 2: High-density dispersal in monoculture

The protocol for experiment 2 was nearly identical to that of experiment 1, except the source patch was inoculated with an aliquot of high-density monoculture (Figure 1); this allowed us to attribute differences in temperature effects on dispersal to density. Five days before conducting experiment 2, we initiated one culture of each species in 100 mL of fresh media and left them to grow at 22 °C. On the day of the experiment, we estimated culture densities in a homogenized 1 mL subsample using a FlowCam (VS Series, Fluid Imaging Technologies, Scarborough, MA) at 40x magnification and a flow rate of 0.3 mL min^-1^. We filled each inoculation patch with 1 mL of stock culture and filled recipient patches with the same volume of protist-free, bacteria-filled media.

As in experiment 1, we recorded start times for each replicate and left them at their respective treatment temperatures for 300 minutes. At the conclusion of the assay, we quickly removed all capillary tubes and added 25 *μ*L of 25% glutaraldehyde to each well to preserve the cells. We quantified the number of preserved individuals in the inoculation and recipient patches of each replicate using the FlowCam.

### 2.4 Experiment 3: Metacommunity colonization rates across temperatures

Each replicate metacommunity was constructed as a 7-patch ring of wells connected by glass capillary tubes. One well was inoculated with a polyculture of all species. This design allowed us to identify when previously empty patches were colonized, allowing quantification of dispersal rates at the leading edge. The ring-shaped configuration allowed us to track colonization in two directions from the patch of origin, and served as insurance against bubbles forming in the capillary tubes and obstructing dispersal (which occurred in a few replicates). Leading edge counts from both directions were averaged to prevent pseudo-replication, resulting in one single estimate of dispersal rate per replicate temperature and species combination.

To test whether thermal performance curve parameters (thermal optimum *T*_opt_; maximum dispersal rate at thermal optimum *d*_max_) for short-term dispersal with little to no population growth (i.e., experiments 1 and 2) could be detected on a longer time scale over which population growth (births, deaths) occurs, we exposed populations to experimental temperatures over multiple generations in the metacommunity landscapes. Wells were sampled at roughly even time intervals (approximately 10-15 hour intervals), generating a time series of species abundances in each patch. We ended the experiment after 60 hours (6-20 generations depending on species and temperature) when all patches had been colonized and new dispersal events could no longer be detected with certainty.

Each inoculation patch was seeded with roughly equal densities of all species (approximately 100 cells mL^-1^). Well plates were then left in incubators set at the same temperatures as before (14 °C, 18 °C, 22 °C, 26 °C, 30 °C, 34 °C), with 6 replicate metacommunities in each temperature treatment (Table 1). We partially sealed the well plates using Parafilm wrap with perforated holes to prevent evaporation while allowing some oxygen exchange and capillary tube insertion.

At each sampling time, we homogenized the contents of the well, transferred a 100 *μ*L subsample onto a sterile microscope slide, and counted all individuals at 40x magnification. We then replaced the 100 *μ*L into its respective well to the best of our ability to reduce temperature-biased population removal, assuming consistent losses.

### 2.5 Replication statement

### 2.6 Analyses

Analyses were performed in R (version 4.5.1, R Core Team 2025).

#### 2.6.1 Fitting thermal response curves for dispersal

We tested for temperature effects on dispersal under no density dependence (Hypothesis 1a) by modelling per capita dispersal rate at low population density, *d*_LD_, as a function of temperature *T* using wait time data from Experiment 1 (Equation 2). Prior to fitting the model, we confirmed that the exponential distribution was a reasonable approximation of the wait time data by visualizing the survival function for each species-by-temperature combination using the Kaplan-Meier estimator (Kaplan and Meier 1958; Figure S1).

Assuming dispersal events reflected a Poisson process, the time to first dispersal follows an exponential distribution with mean waiting time *μ*, where *μ* is modelled as a log-linear function of temperature:

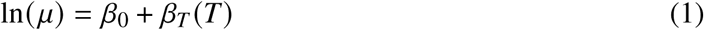

The per-capita dispersal rate is then recovered as:

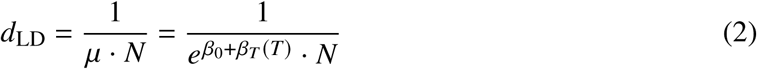

where *N* is the number of individuals inoculated into the source patch and *β_T_* describes the effect of temperature on waiting time. We estimated a separate model for each species using `survreg()` from the `survival R` package (Therneau, 2020). This method allows replicates in which no dispersal event was observed within the 300-minute observation window to contribute information about the lower bound of *d*_LD_ without applying an unobserved value of zero. Confidence intervals on *d*_LD_ were obtained by transforming the standard error of *μ* as 1/(*μ* · *N*) using the delta method (Ver Hoef, 2012), which approximates the variance of a nonlinear function of an estimated parameter as Var (1/*μ*) ≈ Var(*μ*)/*μ*^4^.

We tested for temperature effects on dispersal rates at high population densities (Hypothesis 1b) in high density monoculture (experiment 2) and polyculture (experiment 3). High density dispersal rate (ind ind^-1^ min^-1^) was calculated as the number of individuals in the recipient patch divided by the total population abundance (i.e., sum of individuals in both patches) divided by 300 minutes. Data for *Glaucoma* sp. from experiment 2 were excluded due to poor FlowCam image quality.

We estimated TPCs for high density dispersal rate *d*_HD_ using the *rTPC* and *nls.multstart* R packages (Padfield et al., 2021) to estimate parameters *T*_opt_ (the temperature at which dispersal rate is highest) and *d*_max_ (maximum dispersal rate at *T*_opt_). We performed model selection on four published unimodal TPC models that have strictly positive values (Angilletta 2006; Kontopoulos et al. 2018; Lobry et al. 1991; Lynch and Gabriel 1987; see Table S1 in Supporting Information for the full list). We fitted each model to dispersal rate measurements using a modified Levenberg–Marquardt optimization algorithm with different random combinations of starting parameter values for 1000 iterations, then used the output to estimate the mean corrected Akaike Information Criterion (AICc) score for each TPC model and species. The modified Sharpe-Schoolfield model from Kontopoulos et al. 2018 yielded the lowest average AICc scores for the majority of unimodal TPCs (Table S3) and was subsequently used to estimate TPC parameters of interest in high-density monoculture (experiment 2) and polyculture (experiment 3).

#### 2.6.2 Estimating the strength of density dependence affecting dispersal rates across temperatures

To test H2 (the strength of temperature effects on dispersal vary with population density), we used dispersal rate data from experiments 1 and 2 to estimate density-dependent dispersal rate as a function of population density and temperature (adapted from density-dependent emigration rate in Best et al. 2007 and Altwegg et al. 2013). Dispersal rate from a focal patch to a neighbouring patch varies as a function of temperature *T* and population density *N*:

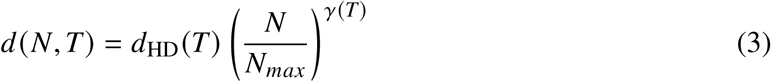

where *d*_HD_(*T*) describes emigration rate at maximum density *N_max_* (analogous to carrying capacity) at a given temperature *T*, *N* is population size at time *t*, and *γ* is a parameter that controls the strength of density dependence. When *N* = *N_max_*, dispersal rate at time *t* is *d*_HD_. When *γ* is positive, dispersal rates increase with population density, and when *γ* = 0, there is no effect of density on dispersal.

We solved for *γ* in equation 4 by substituting population abundances (*N* = abundances in experiment 1, *N_max_* = abundances in experiment 2) and dispersal rates (*d*(*N, T*) = low density dispersal rate, *d*_HD_(*T*) = high density dispersal rate) for each species and temperature.

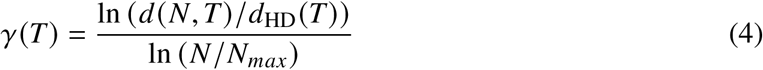

#### 2.6.3 Quantifying dispersal rates in polyculture metacommunities

We quantified leading-edge dispersal rate in the experiment 3 (polyculture metacommunities), *d_p_*, for each species *i* at time point *t* as the proportion of individuals in the leading edge patch post-growth in the preceding leading edge patch since the previous time step:

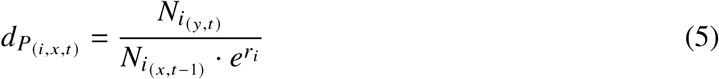

In equation 5, *r_i_* is intrinsic growth rate at the local patch level, x is the leading edge patch at time *t* - 1, and the next patch in the sequence *<u>y</u>* is the leading edge at time *t* (empty or *N_i_* = 0 at time *t* - 1). We used published temperature-dependent intrinsic growth data from Wieczynski et al. (2021), who assayed growth TPCs using the same strains in the present study. Our use of these data assumes that temperature-based differences in *r* measured in single patch monocultures in Wieczynski et al. (2021) approximate the temperature dependence of patch-level growth rates *r_i_* in our experiment. These growth-adjusted dispersal rates were calculated to account for the confounding effects of population growth on observed dispersal rates and used to estimate polyculture dispersal TPCs.

To test H3 (interspecific variation in dispersal TPCs drives differences in metacommunity dynamics across temperatures) we made two comparisons. First, we assessed whether each species’ dispersal TPC in monoculture was preserved in a polyculture metacommunity context by comparing *T*_opt_ and *d*_max_ of high-density monoculture dispersal TPCs with polyculture dispersal TPCs. Second, we assessed whether dispersal TPCs meaningfully change population abundances in multi-patch systems compared to predictions from standard population growth TPCs in single-patch systems. We calculated population growth rate at the whole-metacommunity for species *i*, *r_m,i_*, as:

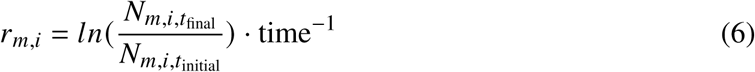

where *N_m_* is the total metacommunity-level abundance of species *i* (Equation 6).

## 3 Results

### 3.1 Hypothesis 1: Temperature affects density-independent and density-dependent dispersal

We tested the hypothesis that temperature affects density-independent dispersal rate (Hypothesis 1a) using survival analysis of time to first dispersal at low population densities (Experiment 1, Equation 1). We found that time to first dispersal decreased significantly with temperature (Figure S1) in *Colpidium striatum* (*β_T_* = -0.05 [95% CI -0.09–0.01], *z* = -2.73, *p* = 0.006), *Glaucoma* sp. (*β_T_* = -0.07 [95% CI -0.11–0.03], *z* = -3.80, *p* < 0.001), and *Paramecium caudatum* (*β_T_* = -0.04 [95% CI -0.08–0.01], *z* = -2.54, *p* = 0.011), translating to increased estimated dispersal rates (*d*_LD_) with temperature in these species (Figure 2a). Temperature had no detectable effect on time to first dispersal in *Paramecium aurelia* (*β_T_* = 0.007 [95% CI -0.03–0.04], *z* = 0.36, *p* = 0.721), *Paramecium bursaria* (*β_T_* = -0.004 [95% CI -0.04–0.04], *z* = -0.18, *p* = 0.861), or *Tetrahymena pyriformis* (*β_T_* = -0.02 [95% CI -0.07–0.03], *z* = -0.78, *p* = 0.433), and predicted *d*_LD_ was comparatively flat across temperatures in these species (Figure 2a). Similarities in cell volume did not translate to similar temperature responses; for example, *Glaucoma* sp. and *Tetrahymena pyriformis* are of roughly comparable size, as are *Paramecium aurelia* and *Paramecium caudatum* (Table S2), yet only one of each pair showed a statistically significant dispersal response to temperature.

**Figure 2:**
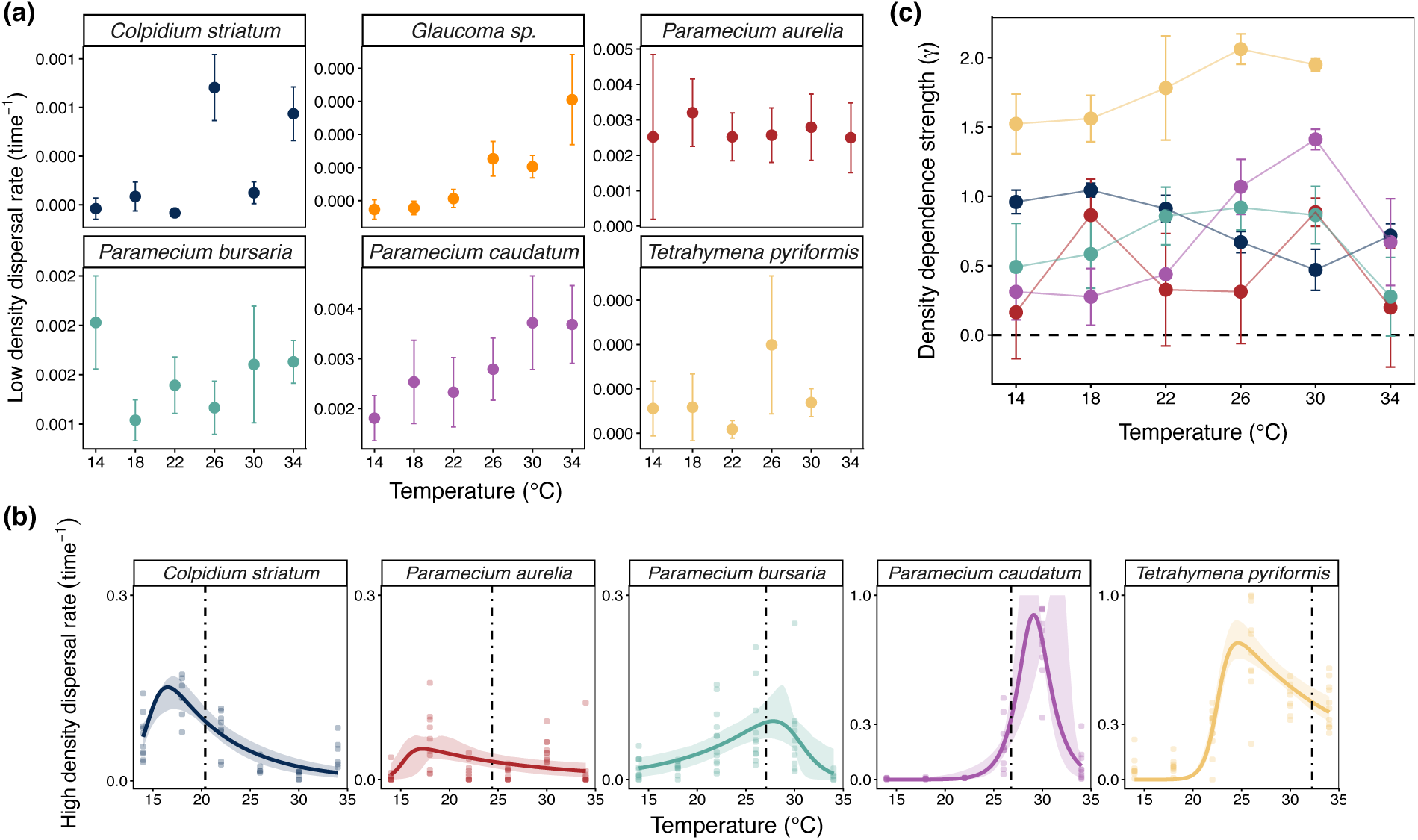
**(a)** Low density dispersal (*d*_LD_) rate versus temperature for six protist species in monoculture. Points indicate the mean estimated per-capita dispersal rate across replicates based on equation 2 with 95% confidence intervals represented as error bars. **(b)** High density dispersal rate (*d*_HD_) versus temperature for five protist species in monoculture. Lines indicate estimated thermal performance curves with bootstrapped 95% confidence intervals for each model fit represented as ribbons. **(c)** Strength of density-dependent effects on dispersal rate (mean *γ*± SD) versus temperature for five protist species. *γ* was estimated using equation 4 fit to data represented in panels (a) and (b).

Consistent with Hypothesis 1b, dispersal rates at high population densities (Experiment 2) showed unimodal responses to temperature in all species (Figure 2b). *Paramecium caudatum* and *Tetrahymena pyriformis* had the highest maximum dispersal rates (89% and 75% of the population dispersing per unit time, respectively) whereas *Paramecium aurelia* had the lowest estimated maximum dispersal rate (10% of the population dispersing per unit time, Figure 2b). Compared to the thermal optimum for population growth, *T*_opt_ for dispersal rate was cold-shifted in *Colpidium striatum*, *Paramecium aurelia*, and *Tetrahymena pyriformis*, slightly warm-shifted in *Paramecium caudatum*, and roughly the same as *T*_opt_ for growth in *Paramecium bursaria* (Figure 2b).

### 3.2 Hypothesis 2: Dispersal rate shows positive density dependence, strength differs among species and temperatures

Density-dependent effects on dispersal rates (*γ*, Equation 3) were nearly always positive, suggesting higher dispersal rates at higher population densities at a given temperature. The strength of density-dependent effects varied across temperatures and species (Hypothesis 2, Figure 2c). In *Paramecium caudatum*, the strength of density dependence showed a clear peak at 30 °C, whereas density dependence was the weakest at this temperature in *Colpidium striatum*. Dispersal rates in *Tetrahymena pyriformis* showed the strongest density dependence among species and temperatures. *Paramecium aurelia* dispersal had the most ambiguous density response, with confidence intervals overlapping 0 at most temperatures; this may reflect uncertainty arising from broadly low dispersal rates and weak temperature effects on dispersal at both low and high population densities (Figure 2).

### 3.3 Hypothesis 3: Metacommunity dynamics reflect interspecific variation in dispersal thermal response curves

In polyculture metacommunities (Experiment 3), differences in dispersal rate TPCs among species were associated with divergent patterns of diversity and abundance across temperatures, consistent with hypothesis 3 (Figure 3, Figure 4). At the first sampling time (*t* = 2), population growth in the inoculation patch (A1) showed a unimodal temperature dependence, with lower growth at 14 °C and 34 °C and highest growth at 26 °C and 30 °C (Figure 3).

**Figure 3:**
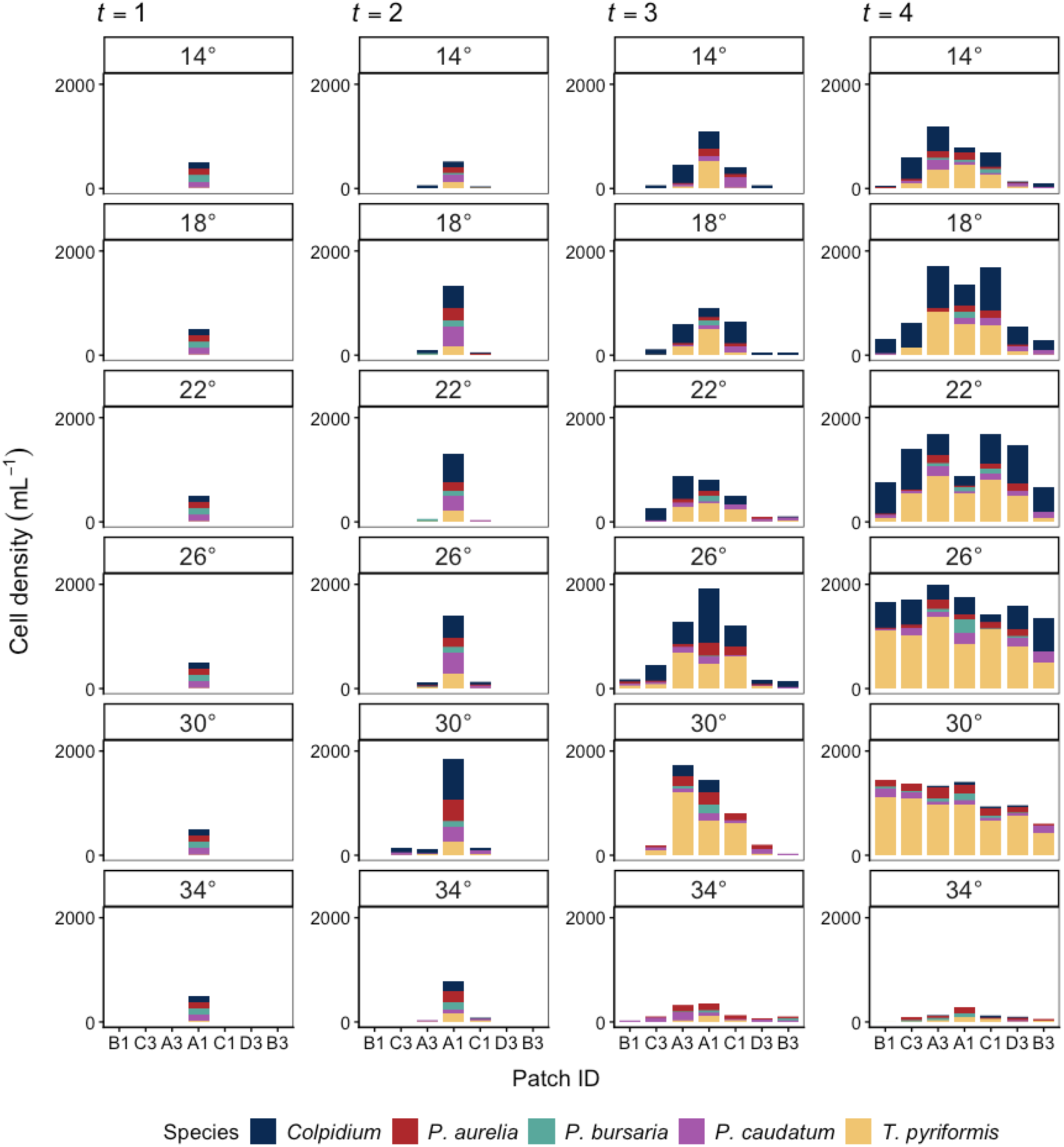
Average species abundances in each patch over the course of the 60-hour metacommunity colonization assay. Average abundances reflect n = 6 replicate metacommunities at each temperature. Panel columns denote the sampling time from left to right, where *t* = 1 is the inoculation time and *t* = 4 is the final sampling time. Patch ID (Well ID) denotes linear position within the metacommunity microcosm, such that Patch ID’s that are adjacent along the horizontal axis were adjacent in the experiment. A1 is the inoculation patch, and protists proliferated outwards from that patch in two directions. A3 and C1, were next in the sequence, and so on until the farthest patches from A1 (B1 and B3) were colonized. Cell densities for numerically dominant *Tetrahymena pyriformis* are divided by 10 for visualization purposes.

**Figure 4:**
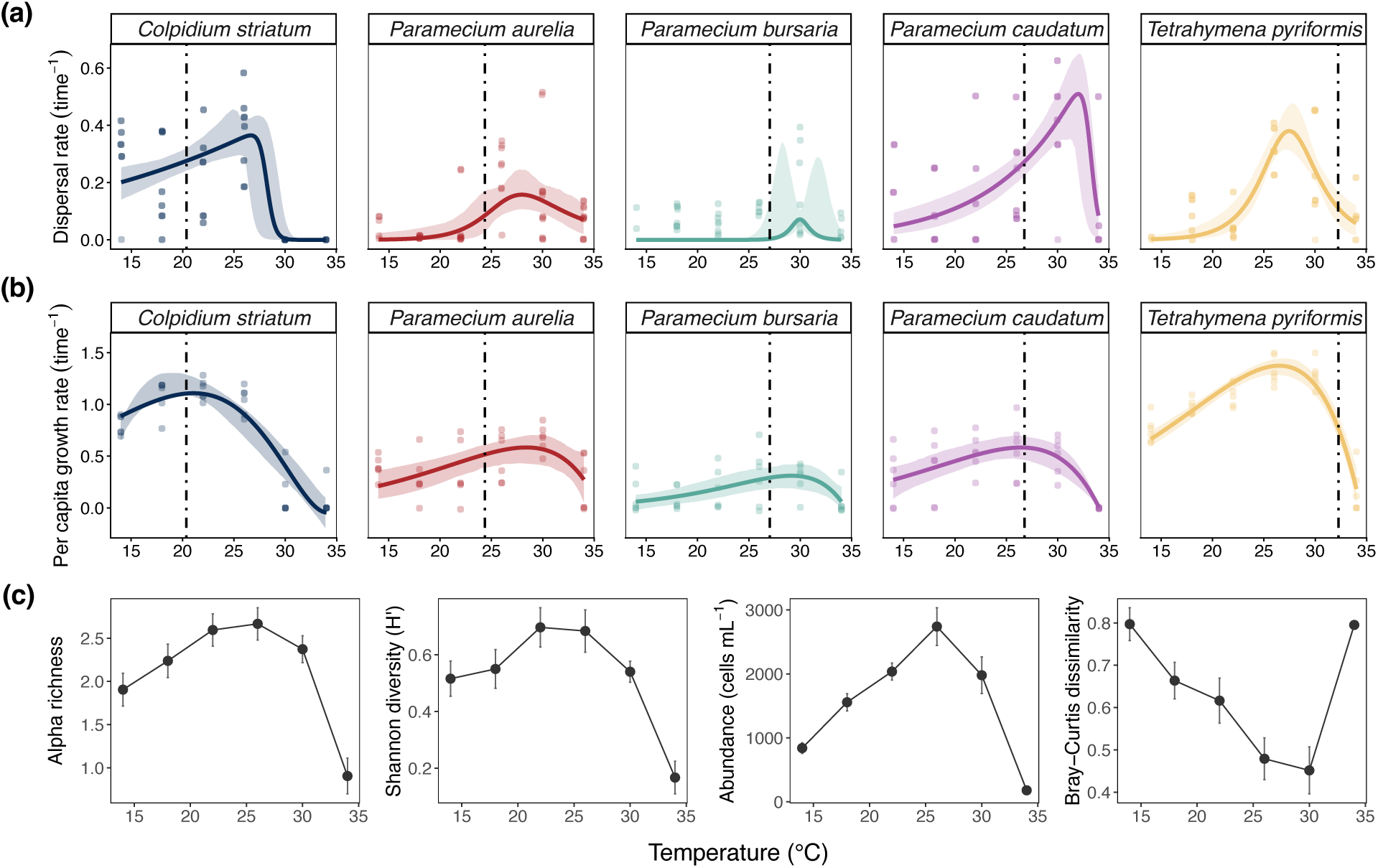
Thermal performance curves for **(a)** leading edge dispersal rates in colonization assay, corrected for effects of population growth, and **(b)** per capita population growth rates at the multi-patch or metapopulation scale (i.e., change in abundance summed across all patches during the metacommunity polyculture assay). Vertical dashed lines indicate species’ thermal optima for intrinsic growth rate in single-patch monoculture (i.e., separate TPC, not shown) for the same strains as published in Wieczynski et al. 2021. **(c)** Summary statistics of local and regional community structure across temperatures in the final time point (*t* = 4).

Leading edge dispersal rate in polyculture, which reflects a time scale exceeding generation time and is corrected for temperature-dependent population growth, varied unimodally with temperature (Figure 4a). All species’ dispersal *T*_opt_ and *d*_max_ were warm-shifted compared to those in high density monoculture (Figure 2b). Polyculture dispersal *T*_opt_ was also warm-shifted compared to the *T*_opt_ for local per capita growth rate (vertical dashed lines) in all species except *Tetrahymena pyriformis*. *Paramecium caudatum* showed the highest of both of these parameters (Figure 4a), as was the case in high density monoculture (Figure 2b). *Paramecium caudatum*’s high dispersal ability at the highest temperatures is reflected in its high relative population abundances in the 30 °C and 34 °C leading edge patches at time point 3 (Figure S2). By the final time point, however, *Tetrahymena pyriformis* was the numerically dominant species at these temperatures.

Patterns of average richness, abundance, and species turnover in the metacommunities resembled unimodal temperature effects (Figure 4c). Species richness and Shannon entropy were highest at 22 °C and 26 °C, while average patch-level abundance was clearly the highest at 26 °C. Bray-Curtis dissimilarity was lowest at 30 °C, reflecting the dominance of *Tetrahymena pyriformis* and absence of other species, particularly in patches farthest from the inoculation patch (Figure S2).

## 4 Discussion

Despite the crucial role of dispersal in population and community dynamics, the effects of temperature on dispersal rates remain poorly understood, particularly in terms of how density-dependence and species-specific thermal performance traits drive regional diversity patterns (Thompson and Gonzalez, 2017). Few studies have explored how temperature-driven changes in dispersal rates might scale up to influence community assembly across spatially structured landscapes. Our study addresses this gap by investigating the temperature dependence of dispersal through a metabolic scaling framework. Measuring dispersal across progressively larger scales of organization allowed us to test whether temperature effects observed in individual dispersal translate into population dispersal rates and metacommunity colonization dynamics. We found that: (1) Density-independent dispersal rate increased with temperature for some species but was invariant for others; (2) High-density dispersal rates exhibited unimodal thermal performance curves, with differences in skewness and thermal optima among species; (3) Population dispersal rates are higher than individual dispersal probabilities due to positive density dependence, and the strength of density dependence differs across temperatures; (4) Temperature-dependent dispersal influences metacommunity-scale population growth in ways that diverge from single-patch monoculture expectations, which, (5) in turn drives differences in local community composition and structure as well as metacommunity dynamics across temperature regimes.

### Temperature dependence of per capita dispersal rate changes with *population density*

We observed some evidence for a monotonic temperature dependence of density-independent dispersal rate at low population densities and on short time scales. Faster swimming speeds were observed at hotter temperatures, but this only translated to a significantly increased dispersal with temperature in half of the study species (Figure 2a). The two-patch arena setup may explain why increased temperature-dependent locomotion did not consistently translate to successful dispersal events. Finding the capillary tube opening was likely influenced in part by swimming speeds but also by the stochastic nature of the protists’ swimming behaviors, such as their frequency and degree of directional changes upon encountering objects (Ishikawa and Kikuchi, 2018; Zhang et al., 2015). Swimming behaviours vary among species, meaning some may have been more likely than others to locate the capillary tube opening in a fixed period of time. Additionally, reduced water viscosity at high temperatures can increase swimming speeds in microbial taxa (Beveridge et al., 2010; Podolsky and Emlet, 1993); we cannot rule out that this non-biological mechanism may have contributed to higher swimming speeds and dispersal at warmer temperatures.

The shift from monotonic low-density dispersal responses to unimodal high-density responses suggests that dispersal TPCs are subject to density dependence. Previous studies show that movement rates in individuals increase monotonically with temperature (Gibert et al., 2016), and this may translate to monotonic dispersal responses at low population densities as observed in experiment 1. However, at high population densities, declines in metabolism-related performance at supra-optimal temperatures may be exacerbated by negative density dependence, which has been well-documented in protists (DeLong et al., 2014; DeLong and Hanson, 2009).

Metabolic scaling theory relates biological processes constrained by aerobic metabolism to the activation energy of aerobic respiration (*E* ↓ 0.65 eV, Gillooly et al. 2001), but biological processes occurring above the level of individuals may have stronger temperature dependencies, reflecting multiple interacting temperature-dependent processes (Anderson-Teixeira et al., 2008; O’Connor et al., 2011). We estimated high thermal sensitivity values for high density population dispersal rates (*E >>* 0.65 eV, Table S4), possibly due to additive effects of temperature on density dependence and movement. This is supported by our finding that the effect of population density on dispersal rate (*γ*, Equation 4) was mostly positive and responded unimodally to temperature for most species (Figure 2c); this observation qualitatively agrees with the hypothesis that per capita metabolic demands correlate with temperature, and thus so too should the strength of density-dependent effects on population processes if resource availability is constant. We acknowledge a limitation in our study arises from the fact that, while starting densities of the bacterial resource community were equal across replicates, we did not quantify temperature differences in bacterial growth at the end of each assay. While beyond the scope of the present study, an interesting future direction may explicitly explore how temperature-dependent trophic effects influence dispersal TPCs.

### 4.1 Short-term dispersal in high density monoculture may depend on both metabolic and behavioural factors

Thermal performance curves for high density dispersal rates varied among species in a manner that did not directly correspond with those for growth rates (Figure 2b). In several species, TPCs for high density dispersal rates were right-skewed, and thermal optima for dispersal were cold-shifted from thermal optima for population growth (Figure 2b). Differences in dispersal rates were not explained by differences in body size, which is consistent with previous findings in ciliates (Beveridge et al., 2010). We hypothesize that dispersal may be influenced by behaviours reflecting thermal preferences that may not correspond to TPCs for growth. A thermal habitat choice experiment with *Tetrahymena thermophila* found that individuals choose to disperse to warmer or cooler patches depending on whether they possess thermal generalist or specialist genotypes (Jacob et al., 2018). Thermal generalists were expected to exhibit random dispersal, as they theoretically should have weaker preferences, however they exhibited non-random dispersal into patches with suboptimal temperatures. Thermal specialists, on the other hand, non-randomly dispersed into patches with optimal temperatures. While our study did not impose temperature differences among neighbouring patches, it is possible that protists sensed their environments relative to their physiological thermal tolerances which influenced non-random emigration behaviours (Jacob et al., 2018). Optimal temperatures for growth may not coincide with optimal temperatures for dispersal if emigration to more favourable environments is not necessary, and dispersal may be elevated at sub- or supra-optimal growth temperatures for the same reason.

### 4.2 Thermal performance curves for dispersal in metacommunities reflect different competition-colonization *trade-offs across temperatures*

Thermal optima for dispersal in polyculture metacommunities (Experiment 3) differed from those measured in monoculture (Experiment 2). Thermal optima for dispersal rate in polyculture were warm-shifted compared to those for growth rates in all but one species (Figure 2b, Figure 4a). A comparison of TPCs for population dispersal rates, single-patch population growth rates, and species-specific whole-metacommunity growth rates suggested potential competition-colonization tradeoffs among species and temperatures. In the metacommunity assay, *Colpidium striatum* had the highest dispersal rates (Figure 4a), often dominating in abundance in the leading edge patches at 26 °C and below (Figure S2). Consistent with this, *Colpidium striatum* exhibited enhanced population growth rates in the metacommunities relative to its expected growth in single-patch monoculture (Figure 4a). *Tetrahymena pyriformis* generally has very high growth rates in monoculture (Wieczynski et al., 2021), and had the highest growth rates of all species in our metacommunity assay. However, its maximum growth rate and growth *T*_opt_ at the whole-metacommunity level were both reduced relative to single-patch monoculture expectations, possibly because its dispersal rates were low (Figure 4).

Having a high dispersal rate at a given temperature may partially offset low growth at that temperature; this may modify a species’ realized thermal performance curve for growth in multi-patch metacommunities relative to those in single-patch monocultures. These results are consistent with previous work examining competition-colonization tradeoffs in ciliates, including several taxa in our study. An experiment found a negative correlation between dispersal and competitive ability in ciliates of different genera and body sizes (Cadotte et al., 2006). If TPCs for dispersal and competitive ability are asymmetrical (i.e., *T*_opt_ for dispersal differs from that of competitive ability) and vary among co-occurring species, then both likely have important effects on changing community composition at different temperatures. Future research could explore whether these competition-colonization trade-offs have a metabolic basis in the form of differential energy allocation between dispersal and resource acquisition that is constrained by temperature.

We observed an interesting anomaly in results from *Paramecium bursaria*, which showed lower dispersal rates and a weak temperature dependence of dispersal in the longer-term polyculture assay. *Paramecium bursaria* is the only ciliate capable of facultative endosymbiosis. While capable of heterotrophy, its ability to compete with other ciliates without its algal endosymbionts was likely diminished due to the bleaching of its endosymbionts (Müller et al., 2012). We observed that *Paramecium bursaria* exhibited more sluggish movements and did not show the same qualitative changes in movement speed with temperature compared to the other species. It is possible that given its ability to acquire carbon from endosymbionts, *Paramecium bursaria* might rely less on movement to acquire food (and in general), translating to lower movement and dispersal rates. Additionally, its thermal optimum for high density dispersal rate in monoculture closely matched its thermal optimum for population growth in monoculture (Figure 2b). *Paramecium bursaria*’s lower reliance on movement for feeding may suggest that its dispersal thermal performance curve more strongly reflects temperature-dependent metabolism, in alignment with classic metabolic theory predictions (Gillooly et al., 2001). Conversely, dispersal thermal responses that deviate from those for growth as in the other species may reflect behavioural adaptations overcoming the pure kinetic effect of temperature. The hypothesis that adaptation and plasticity can overcome kinetic temperature effects on biological rates is known as biochemical adaptation (Kontopoulos et al., 2020) or metabolic compensation (Padfield et al., 2017).

## 5 Conclusion

This study links the effects of temperature on population dispersal rate across time scales and levels of organization to shed light on temperature-dependent metacommunity dynamics. The present findings suggest that population dispersal rates in motile ectotherms are temperature-dependent and influence species’ abundance and persistence at regional scales. Further, thermal performance curves for dispersal vary in several ways: with population density, among species, on different time scales, and in monoculture versus polyculture.

Laboratory microcosms are invaluable tools for robustly testing complex theory in controlled environments (Cadotte et al., 2005), and in a similar vein the general scaling framework presented here for understanding temperature-dependent dispersal can be adapted to other systems. The main mechanisms captured in our three-part experiment—individual dispersal probability, density-dependent dispersal rates, and colonization dynamics in metacommunities—are general features of nearly all species’ dispersal strategies. For example, the transport phase of dispersal in sessile organisms may not depend on locomotion, but is affected by temperature through metabolic constraints on organism development rate and time until settlement (Gillooly et al., 2002; O’Connor et al., 2007). Novel extensions of metabolic theory to include metacommunity dynamics will improve its suitability for addressing common questions in global change ecology (Stark et al., 2025). Our findings yield two main general insights. First, our results support the understanding that temperature effects on diversity and abundance patterns in metacommunities are related to metabolic temperature effects on population dispersal. Second, interspecific differences in dispersal TPCs can introduce variation into realized thermal performance curve shapes for demographic processes at population and community scales. Last and most importantly, ignoring temperature-dependent dispersal may yield an incomplete picture of community dynamics under shifting thermal regimes. Given spatially structured systems experiencing demographic exchange are ubiquitous in nature, the thermal sensitivity of dispersal may prove as consequential as that of any local demographic rate in determining where species persist as climates warm.

## Acknowledgements

We thank Christopher Harley, Michelle Tseng, Rachel Germain, and Amy Angert for feedback on manuscript drafts, and we are grateful to Andrea Yammine, Carling Gerlinsky, Fruin Pow, Sally Jones, Leticia Quieroz, Mairin Netterfield, and Rory Hay for assistance with culture acquisition, maintenance, and running experiments.

## CRediT Author Contribution Statement

**Keila Stark**: Conceptualization, Formal analysis, Investigation, Methodology, Visualization, Writing - original draft. **Ze-Yi Han**: Methodology, Writing - review & editing. **Jean P. Gibert**: Conceptualization, Methodology, Writing - review & editing. **Mary I. O’Connor**: Resources, Supervision, Writing - review & editing.

## Data Availability Statement

Data and code to reproduce analyses are available from https://github.com/keilastark/ protist_dispersal_tpcs and will be published to a permanent repository after publication.

## References

1. Altermatt, F., E. Fronhofer, A. Garnier, A. Giometto, F. Hammes, J. Klecka, D. Legrand, E. Mächler, T. Massie, F. Pennekamp, et al. 2015. Big answers from small worlds: a user’s guide for protist microcosms as a model system in ecology and evolution. methods ecol evol 6: 218–231.

2. Altwegg, R., Y. C. Collingham, B. Erni, and B. Huntley. 2013. Density-dependent dispersal and the speed of range expansions. Diversity and Distributions 19:60–68.

3. Amarasekare, P. 2015. Effects of temperature on consumer–resource interactions. Journal of Animal Ecology 84:665–679.

4. Amarasekare, P. 2024. Temperature-dependent dispersal and ectotherm species’ distributions in a warming world. Journal of Animal Ecology .

5. Anderson-Teixeira, K. J., P. M. Vitousek, and J. H. Brown. 2008. Amplified temperature dependence in ecosystems developing on the lava flows of mauna loa, hawai’i. Proceedings of the National Academy of Sciences 105:228–233.

6. Angilletta, M. J. 2006. Estimating and comparing thermal performance curves. Journal of Thermal Biology 31:541–545.

7. Arancibia, P. A., and P. J. Morin. 2022. Network topology and patch connectivity affect dynamics in experimental and model metapopulations. Journal of Animal Ecology 91:496–505.

8. Barnes, A. D., I. Spey, L. Rohde, U. Brose, and A. I. Dell. 2015. Individual behaviour mediates effects of warming on movement across a fragmented landscape. Functional Ecology 29:1543–1552.

9. Bernhardt, J. R., J. M. Sunday, and M. I. O’Connor. 2018. Metabolic theory and the temperature-size rule explain the temperature dependence of population carrying capacity. The American Naturalist 192:687–697.

10. Best, A., K. Johst, T. Münkemüller, and J. Travis. 2007. Which species will succesfully track climate change? the influence of intraspecific competition and density dependent dispersal on range shifting dynamics. Oikos 116:1531–1539.

11. Bestion, E., J. Clobert, and J. Cote. 2015. Dispersal response to climate change: scaling down to intraspecific variation. Ecology Letters 18:1226–1233.

12. Beveridge, O. S., O. L. Petchey, and S. Humphries. 2010. Mechanisms of temperature-dependent swimming: the importance of physics, physiology and body size in determining protist swimming speed. Journal of Experimental Biology 213:4223–4231.

13. Cadotte, M. W. 2006. Dispersal and species diversity: a meta-analysis. The American Naturalist 167:913–924.

14. Cadotte, M. W., J. A. Drake, and T. Fukami. 2005. Constructing nature: laboratory models as necessary tools for investigating complex ecological communities. Advances in Ecological Research 37:333–353.

15. Cadotte, M. W., D. V. Mai, S. Jantz, M. D. Collins, M. Keele, and J. A. Drake. 2006. On testing the competition-colonization trade-off in a multispecies assemblage. The American Naturalist 168:704–709.

16. Clobert, J., J.-F. Le Galliard, J. Cote, S. Meylan, and M. Massot. 2009. Informed dispersal, heterogeneity in animal dispersal syndromes and the dynamics of spatially structured populations. Ecology letters 12:197–209.

17. Cote, J., E. Bestion, S. Jacob, J. Travis, D. Legrand, and M. Baguette. 2017. Evolution of dispersal strategies and dispersal syndromes in fragmented landscapes. Ecography 40:56–73.

18. DeLong, J. P., T. C. Hanley, and D. A. Vasseur. 2014. Competition and the density dependence of metabolic rates. Journal of Animal Ecology 83:51–58.

19. DeLong, J. P., and D. T. Hanson. 2009. Metabolic rate links density to demography in tetrahymena pyriformis. The ISME journal 3:1396–1401.

20. Doughty, C. E., J. Roman, S. Faurby, A. Wolf, A. Haque, E. S. Bakker, Y. Malhi, J. B. Dunning, and J.-C. Svenning. 2016. Global nutrient transport in a world of giants. Proceedings of the National Academy of Sciences of the United States of America 113:868–873.

21. Eppley, R. W. 1972. Temperature and phytoplankton growth in the sea. Fishery Bulletin 70:1063–1085.

22. Gibert, J. P., M. Chelini, M. F. Rosenthal, and J. P. DeLong. 2016. Crossing regimes of temperature dependence in animal movement. Global Change Biology 22:1722–1736.

23. Gillooly, J. F., J. H. Brown, G. B. West, V. M. Savage, and E. L. Charnov. 2001. Effects of size and temperature on metabolic rate. Science 293:2248–2251.

24. Gillooly, J. F., E. L. Charnov, G. B. West, V. M. Savage, and J. H. Brown. 2002. Effects of size and temperature on developmental time. Nature 417:70–73.

25. Grainger, T. N., and B. Gilbert. 2017. Multi-scale responses to warming in an experimental insect metacommunity. Global Change Biology 23:5151–5163.

26. Hein, A. M., C. Hou, and J. F. Gillooly. 2012. Energetic and biomechanical constraints on animal migration distance: Constraints on animal migration distance. Ecology Letters 15:104–110.

27. Hillaert, J., T. Hovestadt, M. L. Vandegehuchte, and D. Bonte. 2018. Size-dependent movement explains why bigger is better in fragmented landscapes. Ecology and Evolution 8:10754–10767.

28. Hurlbert, A. H., F. Ballantyne, and S. Powell. 2008. Shaking a leg and hot to trot: the effects of body size and temperature on running speed in ants. Ecological Entomology .

29. Ishikawa, T., and K. Kikuchi. 2018. Biomechanics of Tetrahymena escaping from a dead end. Proceedings of the Royal Society B: Biological Sciences 285:20172368.

30. Jacob, S., E. Laurent, B. Haegeman, R. Bertrand, J. G. Prunier, D. Legrand, J. Cote, A. S. Chaine, M. Loreau, J. Clobert, and N. Schtickzelle. 2018. Habitat choice meets thermal specialization: Competition with specialists may drive suboptimal habitat preferences in generalists. Proceedings of the National Academy of Sciences 115:11988–11993.

31. Jetz, W., C. Carbone, J. Fulford, and J. H. Brown. 2004. The scaling of animal space use. Science 306:266–268.

32. Kaplan, E. L., and P. Meier. 1958. Nonparametric estimation from incomplete observations. Journal of the American Statistical Association 53:457–481.

33. Kontopoulos, D.-G., B. García-Carreras, S. Sal, T. P. Smith, and S. Pawar. 2018. Use and misuse of temperature normalisation in meta-analyses of thermal responses of biological traits. PeerJ 6:e4363.

34. Kontopoulos, D.-G., E. van Sebille, M. Lange, G. Yvon-Durocher, T. G. Barraclough, and S. Pawar. 2020. Phytoplankton thermal responses adapt in the absence of hard thermodynamic constraints. Evolution 74:775–790.

35. Lemoine, N. P. 2019. Considering the effects of temperature × nutrient interactions on the thermal response curve of carrying capacity. Ecology 100:e02599.

36. Lobry, J., L. Rosso, and J.-P. Flandrois. 1991. A fortran subroutine for the determination of parameter confidence limits in non-linear models. Binary 3:25.

37. Loreau, M., N. Mouquet, and A. Gonzalez. 2003. Biodiversity as spatial insurance in heterogeneous landscapes. Proceedings of the National Academy of Sciences 100:12765–12770.

38. Lynch, M., and W. Gabriel. 1987. Environmental tolerance. The American Naturalist 129:283–303.

39. Matthysen, E. 2005. Density-dependent dispersal in birds and mammals. Ecography 28:403–416.

40. McNab, B. K. 1963. Bioenergetics and the determination of home range size. The American Naturalist 97:133–140.

41. Mouquet, N., and M. Loreau. 2003. Community patterns in source-sink metacommunities. The american naturalist 162:544–557.

42. Müller, J. P., C. Hauzy, and F. D. Hulot. 2012. Ingredients for protist coexistence: competition, endosymbiosis and a pinch of biochemical interactions. Journal of Animal Ecology 81:222–232.

43. Norberg, J., M. C. Urban, M. Vellend, C. A. Klausmeier, and N. Loeuille. 2012. Eco-evolutionary responses of biodiversity to climate change. Nature climate change 2:747–751.

44. O’Connor, M. I., J. F. Bruno, S. D. Gaines, B. S. Halpern, S. E. Lester, B. P. Kinlan, and J. M. Weiss. 2007. Temperature control of larval dispersal and the implications for marine ecology, evolution, and conservation. Proceedings of the National Academy of Sciences 104:1266–1271.

45. O’Connor, M. I., B. Gilbert, and C. J. Brown. 2011. Theoretical predictions for how temperature affects the dynamics of interacting herbivores and plants. The American Naturalist 178:626–638.

46. Padfield, D., C. Lowe, A. Buckling, R. Ffrench-Constant, Student Research Team, S. Jennings, F. Shelley, J. S. Ólafsson, and G. Yvon-Durocher. 2017. Metabolic compensation constrains the temperature dependence of gross primary production. Ecology Letters 20:1250–1260.

47. Padfield, D., H. O’Sullivan, and S. Pawar. 2021. rTPC and nls.multstart: a new pipeline to fit thermal performance curves in r. Methods in Ecology and Evolution 12:1138–1143.

48. Parain, E. C., S. M. Gray, and L.-F. Bersier. 2019. The effects of temperature and dispersal on species diversity in natural microbial metacommunities. Scientific Reports 9:18286.

49. Pawar, S., A. I. Dell, and Van M. Savage. 2012. Dimensionality of consumer search space drives trophic interaction strengths. Nature 486:485–489.

50. Peters, R. H. 1983. The Ecological Implications of Body Size. 1st ed. Cambridge University Press.

51. Podolsky, R., and R. Emlet. 1993. Separating the effects of temperature and viscosity on swimming and water movement by sand dollar larvae (Dendraster excentricus). Journal of Experimental Biology 176:207–222.

52. R Core Team. 2025. R: A Language and Environment for Statistical Computing. R Foundation for Statistical Computing, Vienna, Austria.

53. Rall, B. C., O. Vucic-Pestic, R. B. Ehnes, M. Emmerson, and U. Brose. 2009. Temperature, predator-prey interaction strength and population stability. Global Change Biology 16:2145–2157.

54. Ritchie, M. G., M. Saarikettu, S. Livingstone, and A. Hoikkala. 2001. Characterization of female preference functions for drosophila montana courtship song and a test of the temperature coupling hypothesis. Evolution 55:721–727.

55. Rocca, J. D., A. Yammine, M. Simonin, and J. P. Gibert. 2022. Protist predation influences the temperature response of bacterial communities. Frontiers in Microbiology 13:847964.

56. Ronce, O., and J. Clobert. 2012. Dispersal syndromes. Dispersal ecology and evolution 155:119–138.

57. Savage, V., J. Gillooly, J. Brown, G. West, and E. Charnov. 2004. Effects of body size and temperature on population growth. The American Naturalist 163:429–441.

58. Schmidt-Nielsen, K. 1972. Locomotion: Energy cost of swimming, flying, and running. Science 177:222–228. Publisher: American Association for the Advancement of Science.

59. Sleigh, M. 1956. Metachronism and frequency of beat in the peristomial cilia of stentor. Journal of Experimental Biology 33:15–28.

60. Stark, K. A., T. Clegg, J. R. Bernhardt, T. N. Grainger, C. P. Kempes, V. Savage, M. I. O’Connor, and S. Pawar. 2025. Toward a more dynamic metabolic theory of ecology to predict climate change effects on biological systems. The American Naturalist 205:285–305.

61. Suárez, D., P. Arribas, E. Jiménez-García, and B. C. Emerson. 2022. Dispersal ability and its consequences for population genetic differentiation and diversification. Proceedings of the Royal Society B 289:20220489.

62. Tamburello, N., I. M. Côté, and N. K. Dulvy. 2015. Energy and the scaling of animal space use. The American Naturalist 186:196–211.

63. Taylor, L. 1963. Analysis of the effect of temperature on insects in flight. The Journal of Animal Ecology pages 99–117.

64. Therneau, T. 2020. A package for survival analysis in r. r package version 3.2-7.

65. Thomas, M. K., C. T. Kremer, C. A. Klausmeier, and E. Litchman. 2012. A global pattern of thermal adaptation in marine phytoplankton. Science 338:1085–1088. Publisher: American Association for the Advancement of Science.

66. Thompson, P. L., B. E. Beisner, and A. Gonzalez. 2015. Warming induces synchrony and destabilizes experimental pond zooplankton metacommunities. Oikos 124:1171–1180.

67. Thompson, P. L., and A. Gonzalez. 2017. Dispersal governs the reorganization of ecological networks under environmental change. Nature Ecology & Evolution 1:0162.

68. Travis, J. M., M. Delgado, G. Bocedi, M. Baguette, K. Bartoń, D. Bonte, I. Boulangeat, J. A. Hodgson, A. Kubisch, V. Penteriani, et al. 2013. Dispersal and species’ responses to climate change. Oikos 122:1532–1540.

69. Travis, J. M., D. J. Murrell, and C. Dytham. 1999. The evolution of density–dependent dispersal. Proceedings of the Royal Society of London Series B: Biological Sciences 266:1837–1842.

70. Tsubaki, Y., Y. Samejima, and M. T. Siva-Jothy. 2010. Damselfly females prefer hot males: higher courtship success in males in sunspots. Behavioral Ecology and Sociobiology 64:1547–1554.

71. Vasseur, D. A. 2020. The impact of temperature on population and community dynamics, pages 243–262. Oxford University Press.

72. Ver Hoef, J. M. 2012. Who invented the delta method? The American Statistician 66:124–127.

73. Wieczynski, D. J., P. Singla, A. Doan, A. Singleton, Z.-Y. Han, S. Votzke, A. Yammine, and J. P. Gibert. 2021. Linking species traits and demography to explain complex temperature responses across levels of organization. Proceedings of the National Academy of Sciences 118:e2104863118.

74. Williams, J. E., and J. L. Blois. 2018. Range shifts in response to past and future climate change: Can climate velocities and species’ dispersal capabilities explain variation in mammalian range shifts? Journal of Biogeography 45:2175–2189.

75. Yamaguchi, R. 2022. Intermediate dispersal hypothesis of species diversity: New insights. Ecological Research 37:301–315.

76. Zhang, P., S. Jana, M. Giarra, P. Vlachos, and S. Jung. 2015. Paramecia swimming in viscous flow. The European Physical Journal Special Topics 224:3199–3210.

