## Supplementary Materials for "Scaling temperature-dependent dispersal rates to metacommunity dynamics: An experimental test"

#### *Supplementary Tables*

**Table S1:** List of candidate thermal response curve models used to fit dispersal rate data from experiment 2 (high density monoculture) and experiment 3 (polyculture). Shorthand model names follow the convention used in the the `rTPC` R package (Padfield et al., 2021).

| Model name | Equation | Reference |
|---|---|---|
| `gaussianmodified_2006` | $d = d_{\max} \cdot exp(-0.5(\frac{\lvert T - T_{\text{opt}} \rvert}{\alpha})^2)$ | Lynch and Gabriel 1987 |
| `lobry_1991` | $d = d_{\max} \cdot (1 - \frac{(T - T_{\text{opt}})^2}{(T - T_{\text{opt}})^2 + T \cdot (T_{\max} + T_{\min} - T) - T_{\max} \cdot T_{\min}}$ | Lobry et al. 1991 |
| `pawar_2018` | $d = \frac{r_{\text{Tref}} \cdot \exp^{-\frac{e}{k}\left(\frac{1}{T+273.15} - \frac{1}{T_{\text{ref}}+273.15}\right)}}{1 + \left(\frac{e}{e_h - e}\right) \cdot \exp^{\frac{e_h}{k}\left(\frac{1}{T_{\text{opt}}+273.15} - \frac{1}{T+273.15}\right)}}$ | Kontopoulos et al. 2018 |
| `weibull_1995` | $d = a \cdot \left(\frac{c-1}{c}\right)^{\frac{1-c}{c}} \left(\frac{T - T_{\text{opt}}}{b} + \left(\frac{c-1}{c}\right)^{\frac{1}{c}}\right)^{c-1} \exp^{-\left(\frac{T - T_{\text{opt}}}{b} + \left(\frac{c-1}{c}\right)^{\frac{1}{c}}\right)^{c}} + \frac{c-1}{c}$ | Angilletta 2006 |

**Table S2:** Inoculation densities for experiment 1 (low-density monoculture) based on roughly equivalent biovolumes of 3-5 individuals of *Paramecium* spp.

| **Species** | **Mean cell biovolume ($\mu$m$^3$)** | **Number of individuals inoculated** |
|---|---|---|
| *Paramecium caudatum* | 287595.3 | 3-5 |
| *Paramecium bursaria* | 196356.3 | 3-5 |
| *Paramecium aurelia* | 185192.4 | 3-5 |
| *Colpidium striatum* | 51087.9 | 30 |
| *Glaucoma* sp. | 17274.3 | 85 |
| *Tetrahymena pyriformis* | 14681.0 | 100 |

**Table S3:** Model selection results for estimated thermal response curves for high density monoculture dispersal rates (experiment 2). The most parsimonious model is shown in bold. Model names correspond with the function name in the `rTPC` R package.

| Species | Model | Sigma | AIC | AICc | BIC | Residual df |
|---|---|---|---|---|---|---|
| *Paramecium caudatum* | **pawar_2018** | 0.10 | -97 | -96 | -87 | 54 |
| | **gaussianmodified_2006** | 0.10 | -97 | -96 | -87 | 54 |
| | **weibull_1995** | 0.10 | -97 | -96 | -87 | 54 |
| | lobry_1991 | 0.12 | -80 | -79 | -69 | 54 |
| *Paramecium aurelia* | pawar_2018 | 0.033 | -234 | -233 | -223 | 56 |
| | gaussianmodified_2006 | 0.032 | -238 | -237 | -227 | 56 |
| | weibull_1995 | 0.033 | -232 | -231 | -221 | 56 |
| | **lobry_1991** | 0.030 | -244 | -243 | -233 | 56 |
| *Paramecium bursaria* | **pawar_2018** | 0.045 | -200 | -199 | -188 | 57 |
| | gaussianmodified_2006 | 0.045 | -198 | -197 | -188 | 57 |
| | weibull_1995 | 0.045 | -199 | -198 | -188 | 57 |
| | **lobry_1991** | 0.045 | -200 | -199 | -190 | 57 |
| *Tetrahymena pyriformis* | **pawar_2018** | 0.15 | -55 | -54 | -45 | 56 |
| | gaussianmodified_2006 | 0.16 | -43 | -42 | -33 | 56 |
| | weibull_1995 | 0.18 | -31 | -30 | -20 | 56 |
| | lobry_1991 | 0.15 | -54 | -52 | -43 | 56 |
| *Colpidium striatum* | **pawar_2018** | 0.033 | -158 | -156 | -149 | 37 |
| | **gaussianmodified_2006** | 0.033 | -158 | -156 | -149 | 37 |
| | weibull_1995 | 0.033 | -157 | -155 | -149 | 37 |
| | lobry_1991 | 0.042 | -137 | -136 | -129 | 37 |

**Table S4:** Estimated thermal response curve parameters for high density dispersal rate in monoculture (experiment 2). Parameters correspond with thermal response curves in main text figure 2a.

| Species | $d_{max}$ | $T_{opt}$ | $CT_{min}$ | $CT_{max}$ | E |
|---|---|---|---|---|---|
| *Paramecium caudatum* | 0.89 | 29.10 | 24.05 | 34.83 | 5.73 |
| *Paramecium aurelia* | 0.05 | 17.25 | 13.08 | 58.38 | 3.44 |
| *Paramecium bursaria* | 0.10 | 27.80 | 4.46 | 35.19 | 0.92 |
| *Tetrahymena pyriformis* | 0.75 | 24.63 | 20.30 | 64.75 | 1.92 |
| *Colpidium striatum* | 0.15 | 16.51 | 11.35 | 38.06 | 6.80 |

**Table S5:** Estimated thermal response curve parameters for different species in polyculture (experiment 3). Parameters correspond with thermal response curves in main text figure 4a.

| **Species** | $\mathbf{d_{max}}$ | $\mathbf{T_{opt}}$ | $\mathbf{CT_{min}}$ | $\mathbf{CT_{max}}$ | **E** |
|---|---|---|---|---|---|
| *Paramecium caudatum* | 0.32 | 32.64 | 15.31 | 35.11 | 1.36 |
| *Paramecium aurelia* | 0.29 | 29.64 | 24.65 | 32.03 | 7.38 |
| *Paramecium bursaria* | 0.032 | 20.64 | 14.83 | 35.30 | 0.15 |
| *Tetrahymena pyriformis* | 0.029 | 28.32 | -4.41 | 30.93 | 0.65 |
| *Colpidium striatum* | 0.18 | 25.89 | 2.24 | 33.42 | 0.96 |

### *Supplementary Figures*

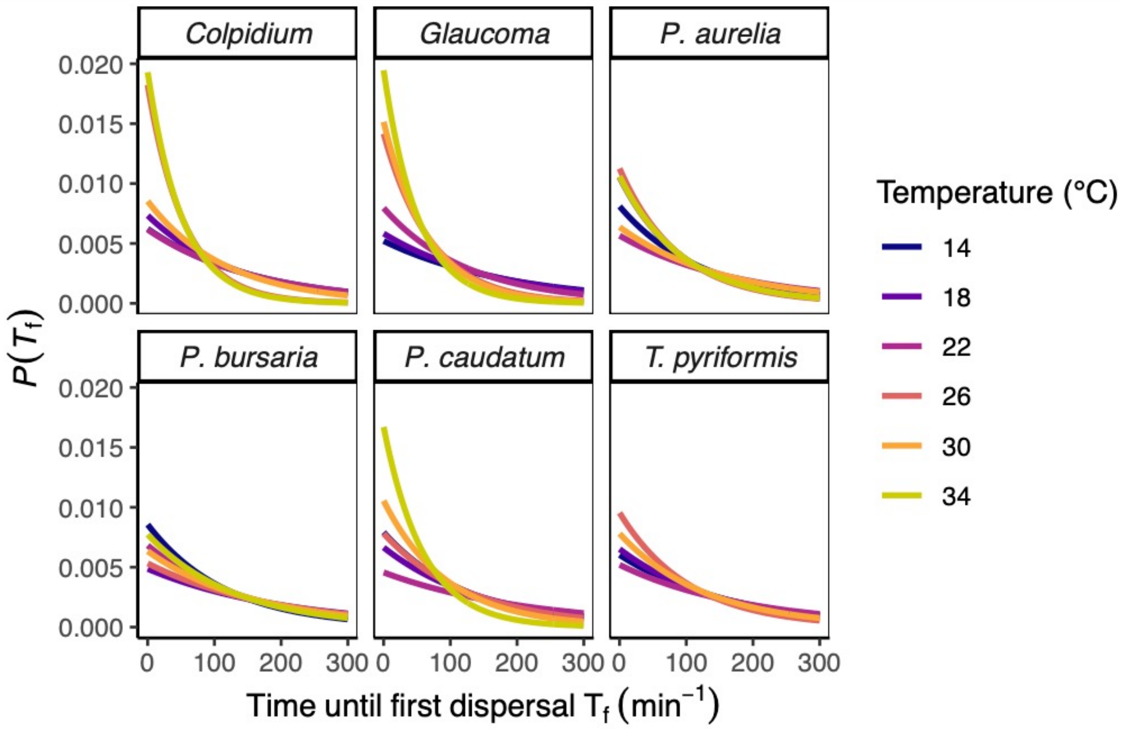


**Figure S1:** Probability density function for the exponential distribution, representing low-density dispersal probability (experiment 1) as a function of wait times at different temperatures. Steeper curves indicate a shorter time until first dispersal (i.e., higher exponential distribution rate parameter).

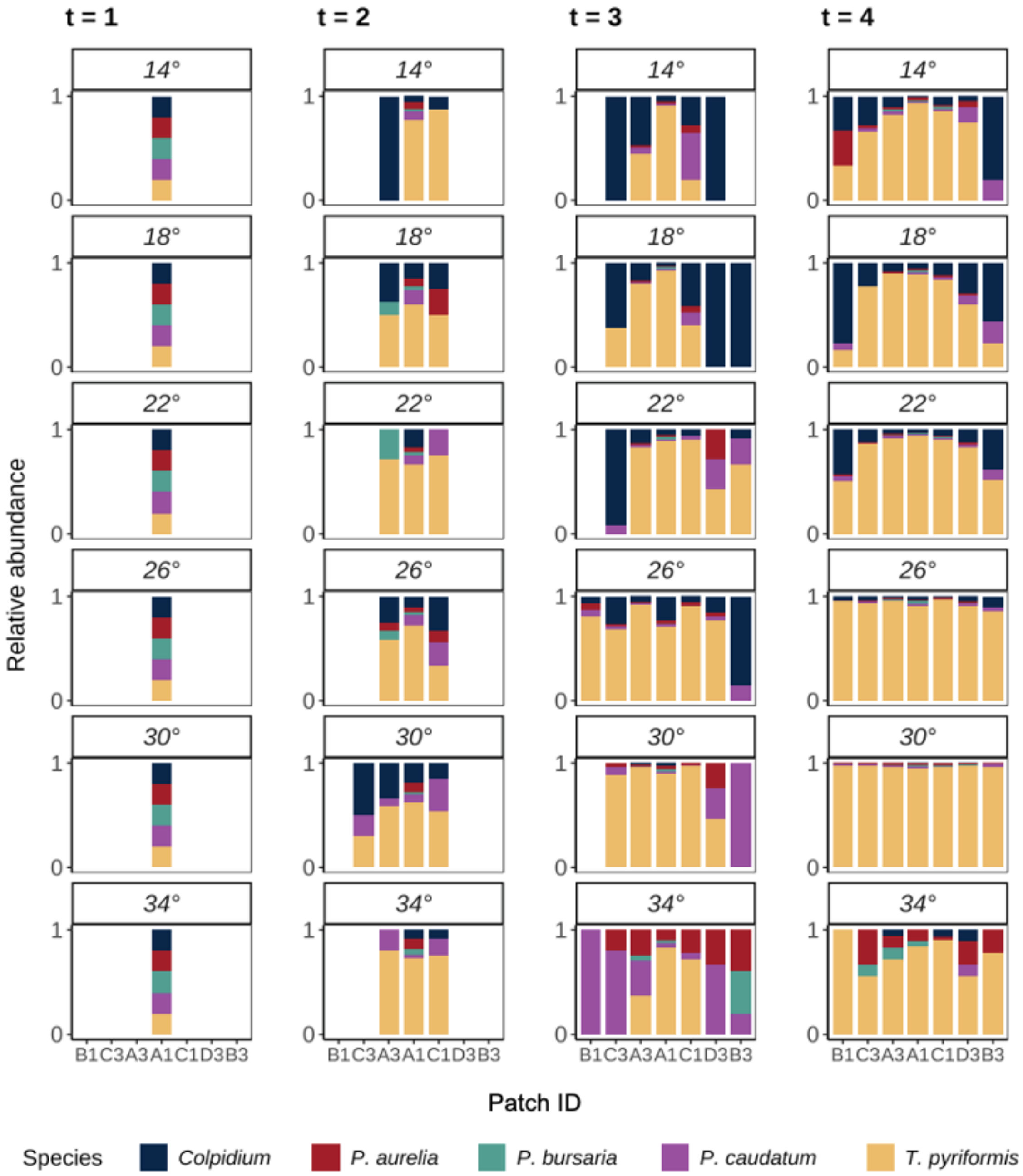


**Figure S2:** Average proportional abundances in each patch over the course of the 60-hour metacommunity colonization assay. Average abundances reflect n = 6 replicate metacommunities at each temperature. Panel columns denote the sampling time from left to right, where $t$ = 1 is the inoculation time and $t$ = 4 is the final sampling time. Patch ID (Well ID) denotes linear position within the metacommunity microcosm, such that Patch ID's that are adjacent along the horizontal axis were adjacent in the experiment. A1 is the inoculation patch, and protists proliferated outwards from that patch in two directions. A3 and C1, were next in the sequence, and so on until the farthest patches from A1 (B1 and B3) were colonized.
